# Nuclear localization of TORC1 and cellular co-localization of TORC1 and c-Fos in the visceral neuraxis after systemic LiCl injection

**DOI:** 10.1101/259812

**Authors:** Adam Kimbrough, Thomas A. Houpt

**Author notes:** Correspondence: Thomas A. Houpt, Department of Biological Science, 3010 King Building, The Florida State University, Tallahassee, FL 32306-4295, USA.

## Abstract

Cyclic–AMP response element binding protein (CREB)-mediated gene expression is critical for the processing of visceral information, including learning about visceral stimuli (e.g. conditioned taste aversion). However, CREB requires additional co-factors to induce gene expression, including transducer of regulated CREB-activity (TORC). The nuclear localization of TORC1 has not been examined previously in the visceral neuraxis. c-Fos, a widely-used marker of activity, is induced by visceral stimulation. If CREB-mediated gene transcription is necessary for c-Fos induction in the visceral neuraxis after systemic LiCl injection, then TORC should be active following stimulation, and co-localized with c-Fos in activated neurons. We examined nuclear (activated) TORC1 in the visceral neuraxis at 30, 60, and 180 minutes after LiCl, as compared to c-Fos induction. Consistent with previous studies we found increases in c-Fos 60 min after LiCl in all regions. Nuclear TORC1 was also increased in the area postrema, NTS, and paraventricular nucleus after LiCl. Surprisingly, in the central amygdala, TORC1 was deactivated after LiCl, such that almost all TORC1 was cytoplasmic, whereas control rats had almost all nuclear TORC1. Fluorescent double-labeling of TORC1 and c-Fos after LiCl found that cellular co-localization of c-Fos and TORC1 was very low across the visceral neuraxis. Nuclear TORC1 expression reveals a population of cells in the visceral neuraxis, independent of c-Fos-positive cells, that are regulated by LiCl. TORC1, and in turn CREB-mediated gene transcription, may not be necessary for c-Fos induction in the visceral neuraxis after LiCl.

## 1. Introduction

The cyclic-AMP (cAMP)-Protein Kinase A (PKA)-cAMP response element-binding protein (CREB) pathway mediates much gene expression in the brain following visceral stimulation, and during acquisition in many forms of learning and memory, including conditioned taste aversion (CTA) learning. Within the nucleus, phosphorylated CREB (pCREB), a substrate of PKA, mediates critical gene transcription by binding to CREB response elements (CREs) within the promoter of target genes (Mayr and Montminy, 2001). Thus, CREB-inducible gene products are candidate mediators of visceral responses, and may contribute to long-term consolidation of visceral CTA learning.

Visceral neuraxis activation by interoceptive stimuli is accompanied by CREB-inducible gene expression, and induction of genes such as the immediate-early gene c-Fos. For example, lithium chloride (LiCl), a commonly used toxic US in CTA protocols, induces c-Fos expression along the visceral neuraxis, i.e., in the nucleus of the solitary tract (NTS), parabrachial nucleus (PBN), supraoptic nucleus (SON), paraventricular nucleus (PVN), and central amygdala (CeA) (Gu et al., 1993; Houpt et al., 1994; Koehnle and Rinaman, 2007; Lamprecht and Dudai, 1995; Nunnink et al., 2007; Sakai and Yamamoto, 1997; Spencer and Houpt, 2001; St Andre et al., 2007; Swank and Bernstein, 1994; Yamamoto et al., 1992; Yamamoto et al., 1997). LiCl-induced c-Fos is dependent on both neural and humoral inputs to the visceral neuraxis. Lesions of the area postrema (AP) or vagus nerve attenuate or eliminate c-Fos after LiCl in some brain regions (Spencer et al., 2012; Yamamoto et al., 1992). Within the visceral neuraxis, transynaptic activation after LiCl injection depends on the release of neurotransmitters such as glutamate and GLP-1 from hindbrain projections onto forebrain sites (Kinzig et al., 2002; Seeley et al., 2000).

c-Fos is a candidate target gene of CREB during visceral activation and CTA acquisition. c-Fos transcription can be initiated by phosphorylated CREB (pCREB) binding to CRE sites in the c-Fos promoter (Herdegen and Leah, 1998; Mayr and Montminy, 2001). Furthermore, CREB and pCREB are present in brain regions where LiCl induces c-Fos (Houpt, 1997; Lamprecht et al., 1997; Oberbeck et al., 2010; Shiromani et al., 1995). However, pCREB alone may not be sufficient to induce c-Fos expression. For example, many regions show high levels of pCREB in the basal state without basal c-Fos expression (Houpt, 1997; Lamprecht et al., 1997; Oberbeck et al., 2010; Shiromani et al., 1995). Thus, the immunohistological data alone suggests that other transcription factors, in addition to CREB, contribute to c-Fos expression.

One class of transcription factors that might also contribute to LiCl-induced c-Fos is transducer of Regulated CREB activity (TORC) proteins, which have been recently identified as contributors to CREB-mediated gene expression. There are 3 subtypes of TORC, all of which are expressed in the brain, with TORC1 being the most abundant (Watts et al. 2011). TORC is sequestered in the cytoplasm in a phosphorylated state by protein 14-3-3 as shown by immunoprecipitation in HEK cells (Screaton et al., 2004). In the presence of calcium, TORC is dephosphorylated by calcineurin (Bittinger et al., 2004; Kingsbury et al., 2007; Screaton et al., 2004; Spencer and Weiser, 2010) and then translocates to the nucleus, where it binds to CREB as a required co-factor to facilitate gene transcription (Bittinger et al., 2004; Conkright et al., 2003a; Conkright et al., 2003b; Iourgenko et al., 2003).

TORC proteins have been shown to contribute to CREB-mediated gene transcription in several systems, such as gluconeogenesis in the liver, ischemia in cortex, and long-term potentiation in the hippocampus (Jansson et al., 2008; Sasaki et al., 2011; Zhou et al., 2006). In the visceral neuraxis, nuclear TORC2 in the PVN increases following restraint stress, which correlates with increased corticotropin releasing hormone (CRH) mRNA (Liu et al., 2011).

So, CRE-mediated gene transcription requires both phosphorylation of CREB within the nucleus and translocation of TORC from the cytoplasm to the nucleus. Thus, if LiCl injection induces CREB-mediated gene transcription in the visceral neuraxis leading to c-Fos induction, then i) nuclear TORC should be present, e.g., in the site of c-Fos expression, and ii) TORC should be increased following LiCl injection and preceding c-Fos induction. However, TORC has not been examined after acute LiCl injection, nor has TORC been co-localized in the nucleus of cells expressing c-Fos after acute LiCl.

Therefore, to establish the relation of TORC expression with c-Fos induced by LiCl in the visceral neuraxis,

1) We examined changes in nuclear TORC1 immunoreactivity at 30, 60, and 180 min after LiCl injection. TORC1 is one of 3 isoforms of TORC and is the most abundant isoform in the brain (Li et al., 2009; Watts et al., 2011). A previous report by Li et al. (2009) showed that nuclear TORC1 is elevated at 30 min after stimulation and reaches maximal nuclear levels at 60 min in cultured cortical neurons. Thus, TORC1 nuclear localization was predicted at 30 min so as to precede c-Fos protein induction at 60 min.

2) We examined nuclear co-localization of c-Fos with TORC1 at 60 min after LiCl, when c-Fos protein levels are maximal (Spencer and Houpt, 2001), by fluorescent double-labeled immunohistochemistry.

## 2. Results

### 2.1 Experiment 1: Time course of LiCl-induced nuclear TORC1 expression

#### 2.1.1 Basal Levels of TORC1

Rats were either perfused immediately with no treatment or 30 min after NaCl injection, and then brains were processed for immunohistochemistry to establish basal levels of TORC1 nuclear localization and the effects of an injection procedure in the visceral neuraxis. Nuclear TORC1 in the AP, NTS, PBN, SON, PVN, and CeA was quantified.

Basal levels of nuclear TORC1 were high in all brain regions examined. In the AP, NTS, PBN, SON, and CeA there was no difference between saline-injected and no-treatment groups in number of cells expressing nuclear TORC1 (see Figure 1). However, in the PVN the no-treatment group had a significantly greater number of nuclear TORC1 cells than the 30-min NaCl group (p<.05).

**Figure 1.**
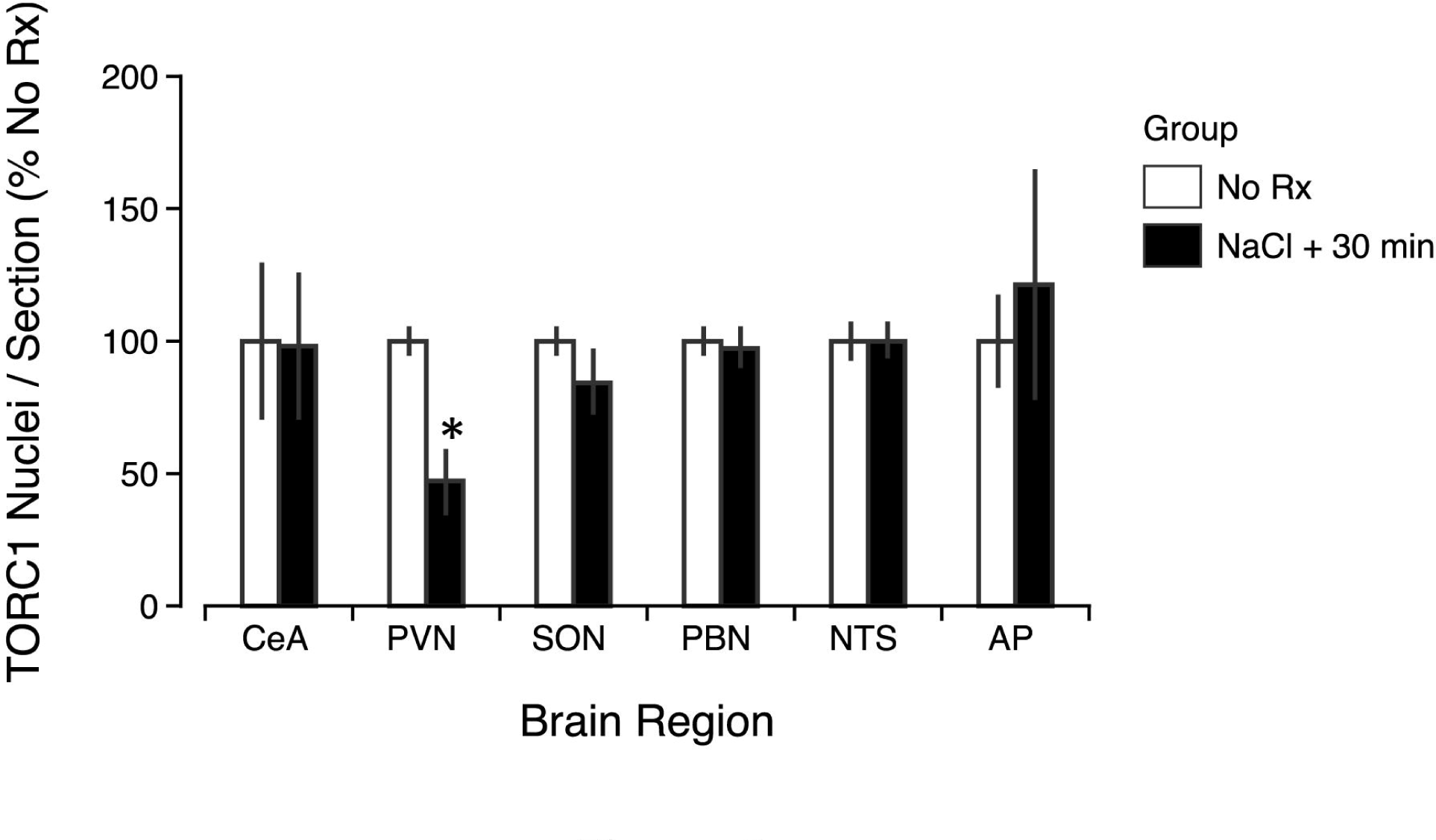
Basal TORC1 levels in the visceral neuraxis. Cell counts for nuclear TORC1 in the AP, NTS, PBN, PVN, SON, and CeA after no treatment or 30 min after NaCl injection. No differences were seen between groups, except for in the PVN where NaCl-injected rats had significantly lower nuclear TORC1 than rats receiving no treatment. * p<.05 significantly different from no-treatment group.

**Figure 2.**
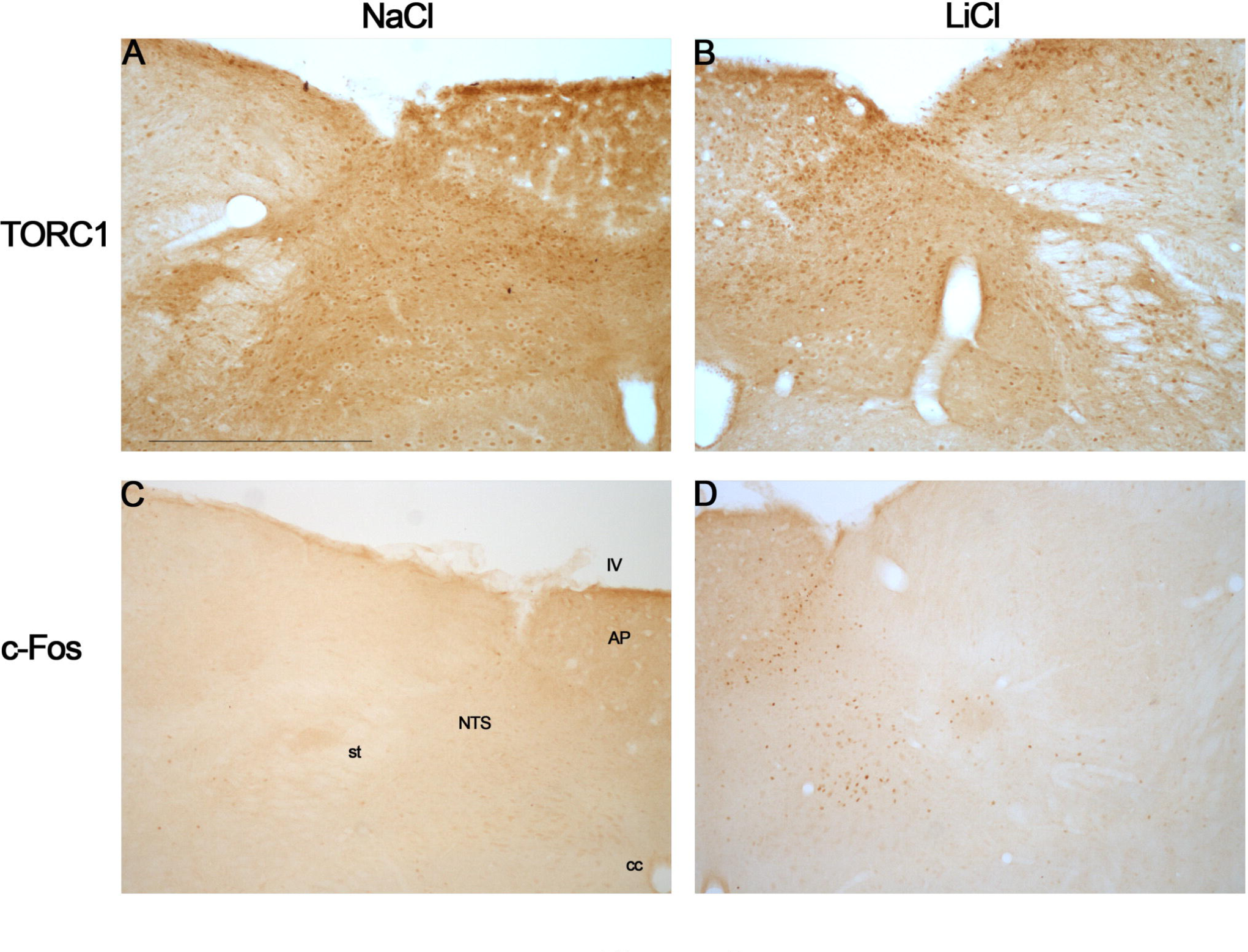
Photomicrographs of the AP and NTS showing TORC1 immunohistochemistry after injections of NaCl (A) and LiCl (B) and c-Fos immunohistochemistry after injections of NaCl (C) and LiCl (D). st, solitary tract; NTS, nucleus of the solitary tract; cc, central canal; AP, area postrema; IV, fourth ventricle. Scale bar 300 µm.

In order to examine the effects of LiCl injection on nuclear TORC1 localization in the visceral neuraxis, rats were injected with LiCl or NaCl and perfused at 30, 60 or 180 min after injection. Rats were processed for chromogenic immunohistochemistry of TORC1 (30, 60 and 180 min) and c-Fos (60 min).

#### 2.1.2 TORC1 and c-Fos induction after LiCl Injection

Nuclear c-Fos was significantly elevated (p<.05) in LiCl-injected rats compared to NaCl-injected rats at 60 min in all regions of the visceral neuraxis examined (the AP, NTS, PBN, SON, PVN, and CeA; see Figures 3 and 7). The c-Fos labeling was clustered and very nuclear and punctate.

**Figure 3.**
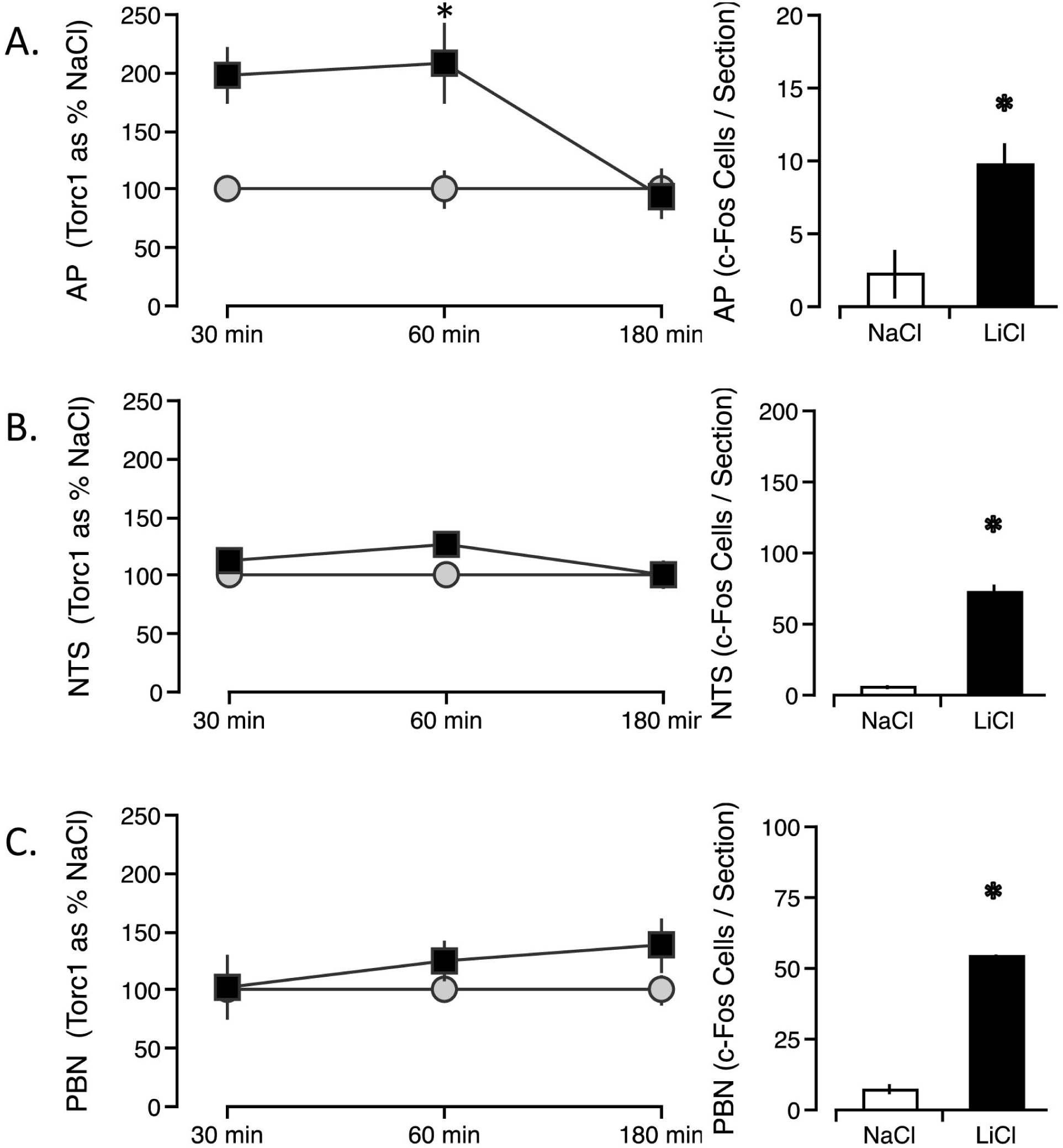
Time course of nuclear TORC1 and c-Fos in the hindbrain of the visceral neuraxis following i.p. LiCl injection (black symbols) or NaCl injection (white symbols). Nuclear TORC1 was significantly increased in the AP (A) and NTS (B) in LiCl-injected rats when compared to NaCl-injected rats. No significant difference in nuclear TORC1 was seen in the PBN (C). c-Fos was significantly elevated at 60 min in all LiCl-injected rats compared to NaCl-injected rats (A,B,C). * p<.05 significantly different from NaCl-injected.

**Figure 4.**
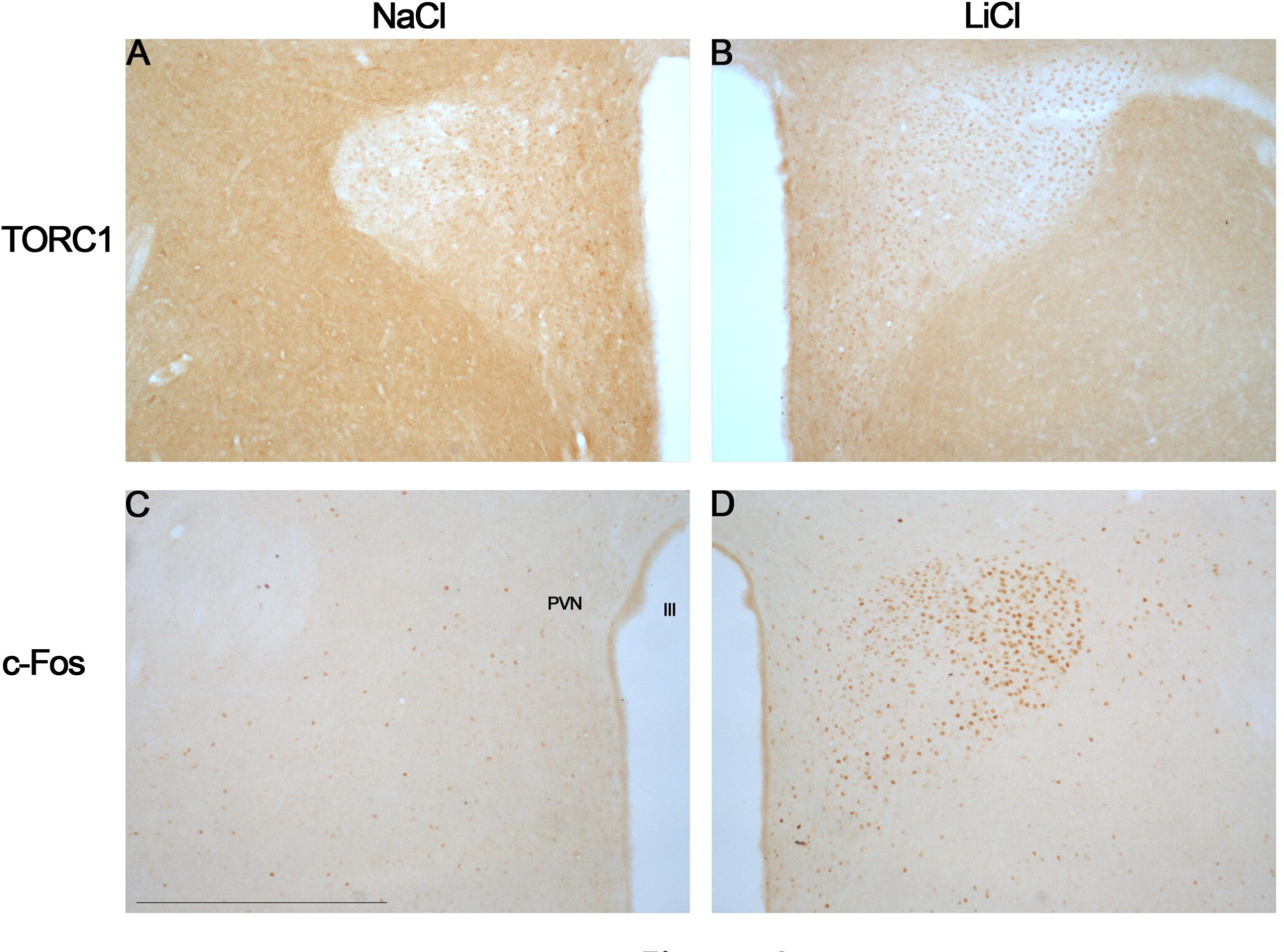
Photomicrographs of the PVN showing TORC1 immunohistochemistry after injections of NaCl (A) and LiCl (B) and c-Fos immunohistochemistry after injections of NaCl (C) and LiCl (D). iii, third ventricle; PVN, paraventricular nucleus. Scale bar 300 µm.

**Figure 5.**
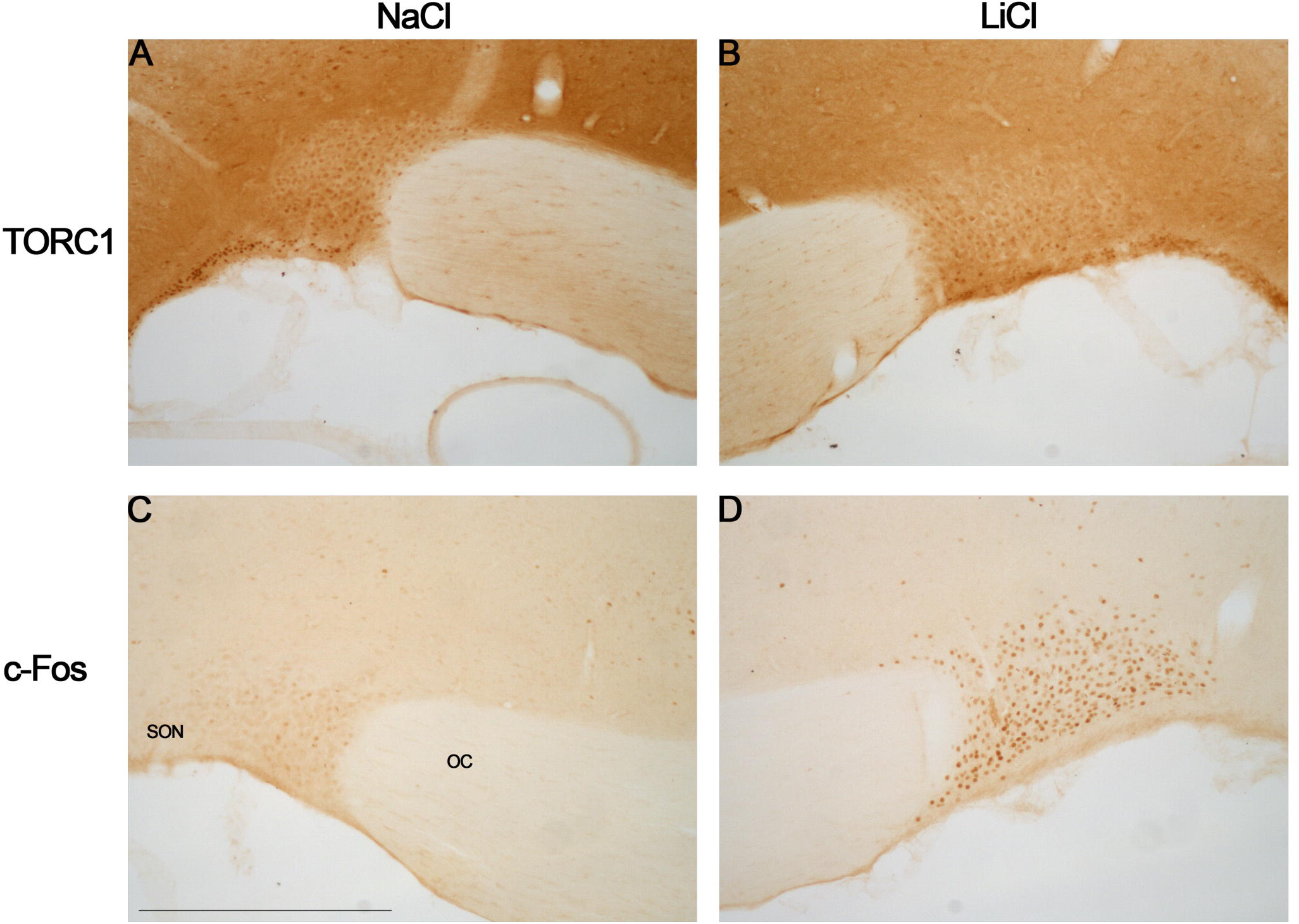
Photomicrographs of the SON showing TORC1 immunohistochemistry after injections of NaCl (A) and LiCl (B) and c-Fos immunohistochemistry after injections of NaCl (C) and LiCl (D). oc, optic chiasm; SON, supraoptic nucleus. Scale bar 300 µm.

**Figure 6.**
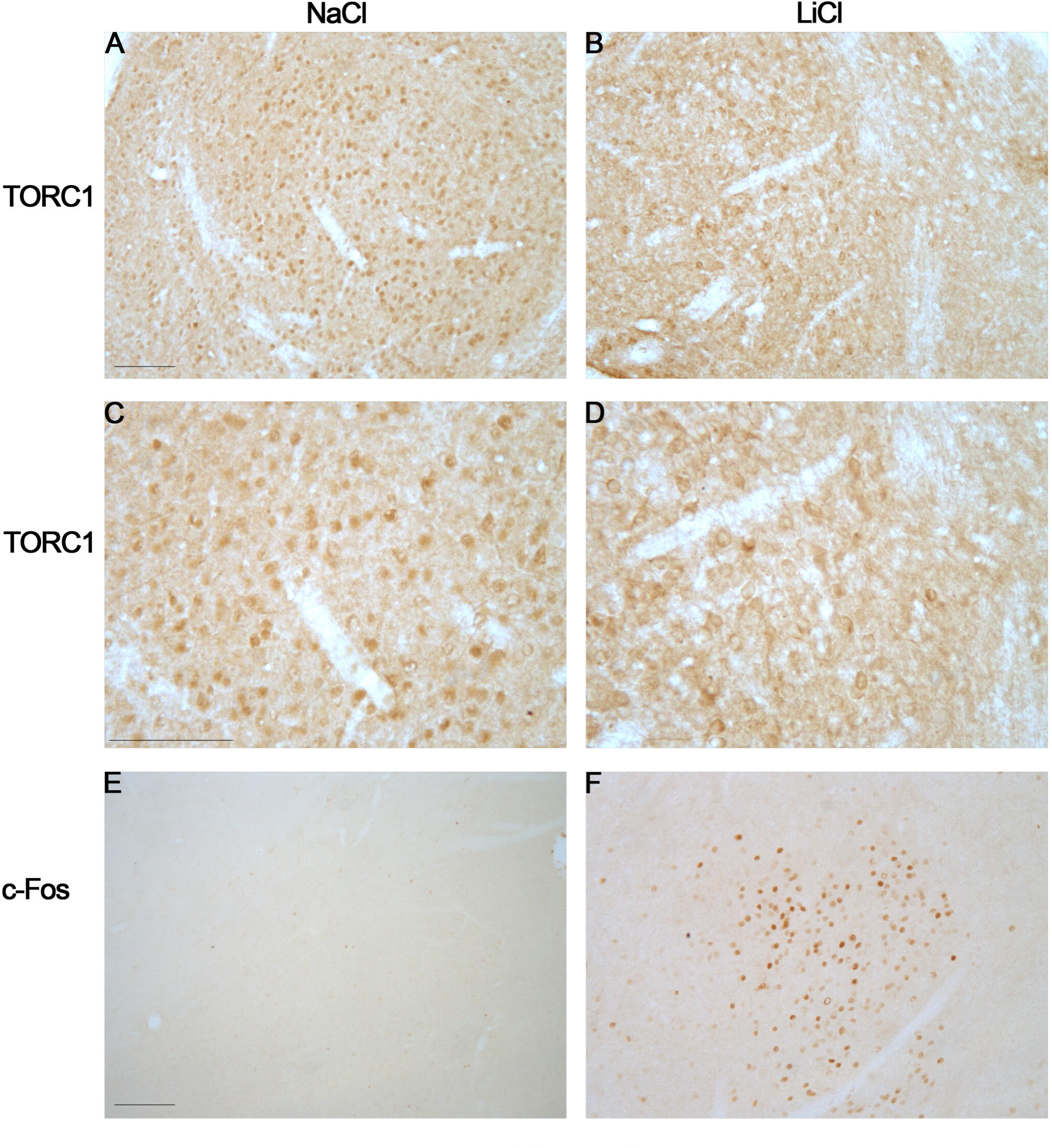
Photomicrographs of the CeA showing TORC1 and c-Fos immunohistochemistry after LiCl injection. TORC1 in the CeA after injections of NaCl (A) and LiCl (B) and higher magnification NaCl (C) and LiCl (D). c-Fos in the CeA after injections of NaCl (E) and LiCl (F). Scale bar 70 µm.

**Figure 7.**
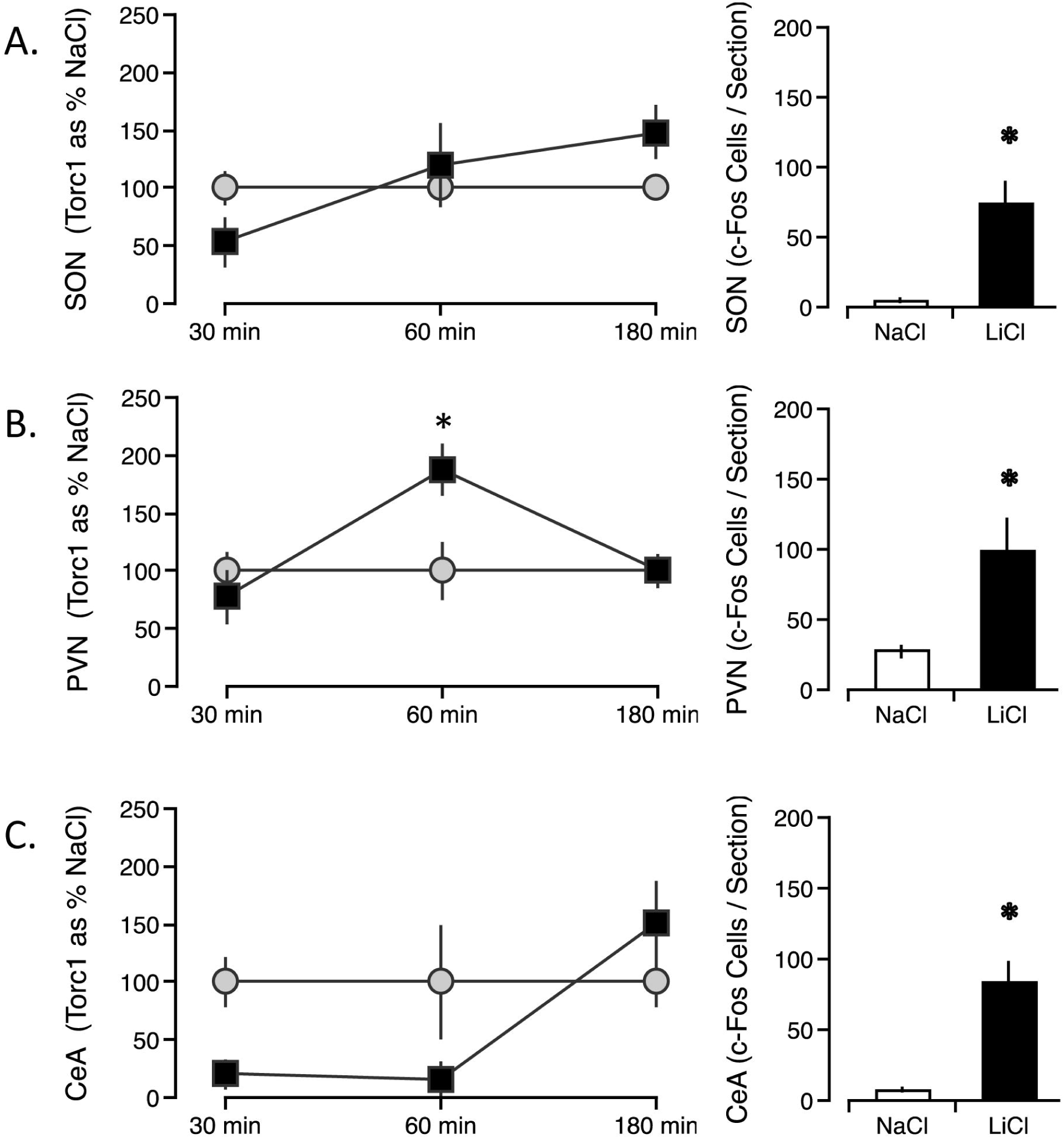
Time course of nuclear TORC1 and c-Fos induction in the forebrain of the visceral neuraxis following i.p. LiCl injection (black symbols) or NaCl injection (white symbols). Nuclear TORC1 was significantly increased in the PVN (A) in LiCl-injected rats when compared to NaCl-injected rats. Nuclear TORC1 was significantly reduced in the CeA (B) of LiCl-injected rats compared to NaCl injected-rats. No significant difference in nuclear TORC1 was seen in the SON (C). c-Fos was significantly elevated at 60 min in all LiCl-injected rats compared to NaCl-injected rats (A,B,C). * p<.05 significantly different from NaCl-injected.

**Figure 8.**
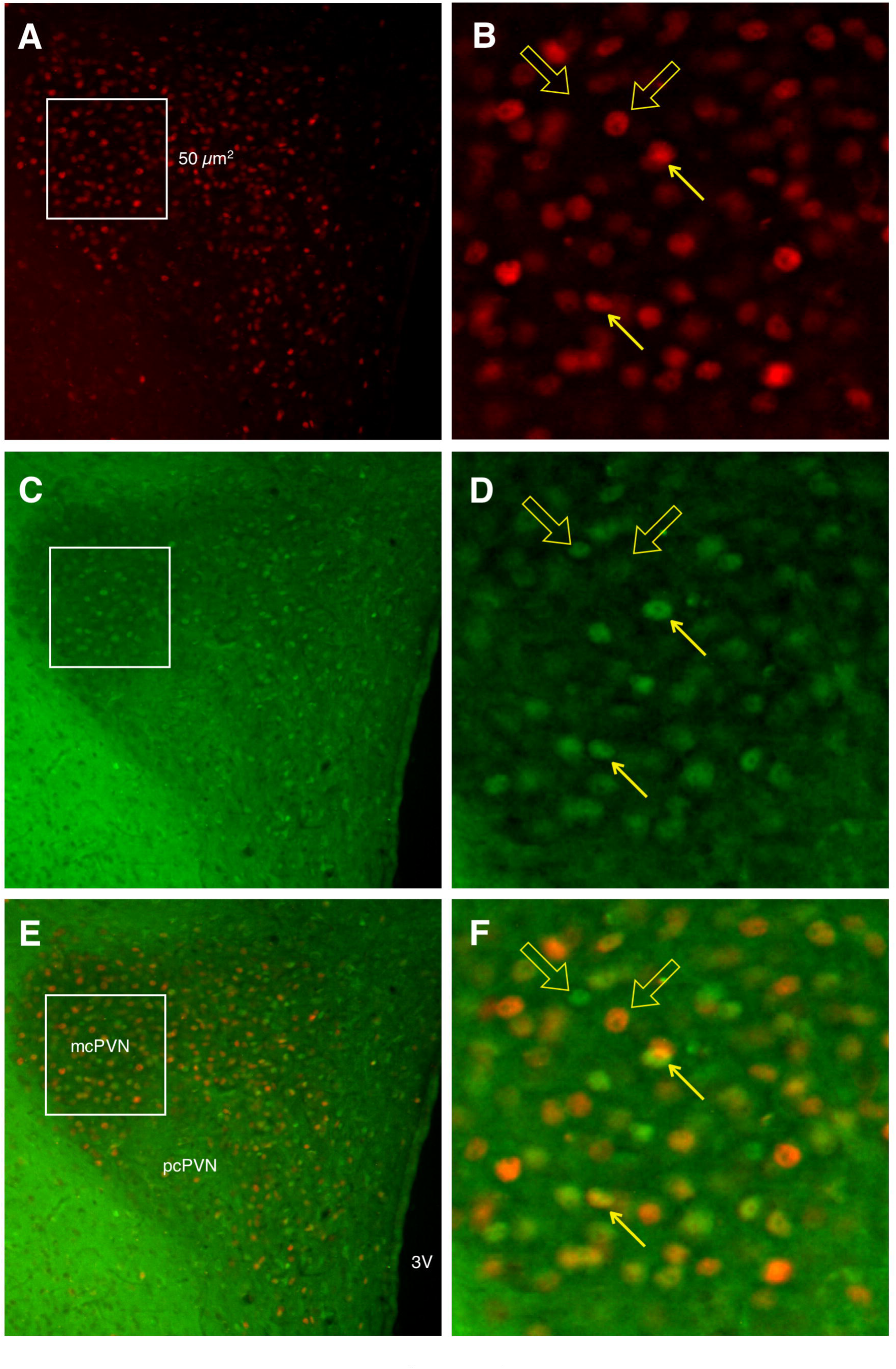
Photomicrographs of fluorescent immunohistochemistry of TORC1 and c-Fos in the PVN 60 min after LiCl injection. Red nuclear c-Fos labeling (A) and magnified (B). Green TORC1 staining (C) and magnified (D). Merged c-Fos and TORC1 labeling (E) and magnified (F). 3V, third ventricle; mcPVN, magnocellular paraventricular nucleus; pcPVN, parvocellular paraventricular nucleus. Scale bar 50 µm.

**Figure 9.**
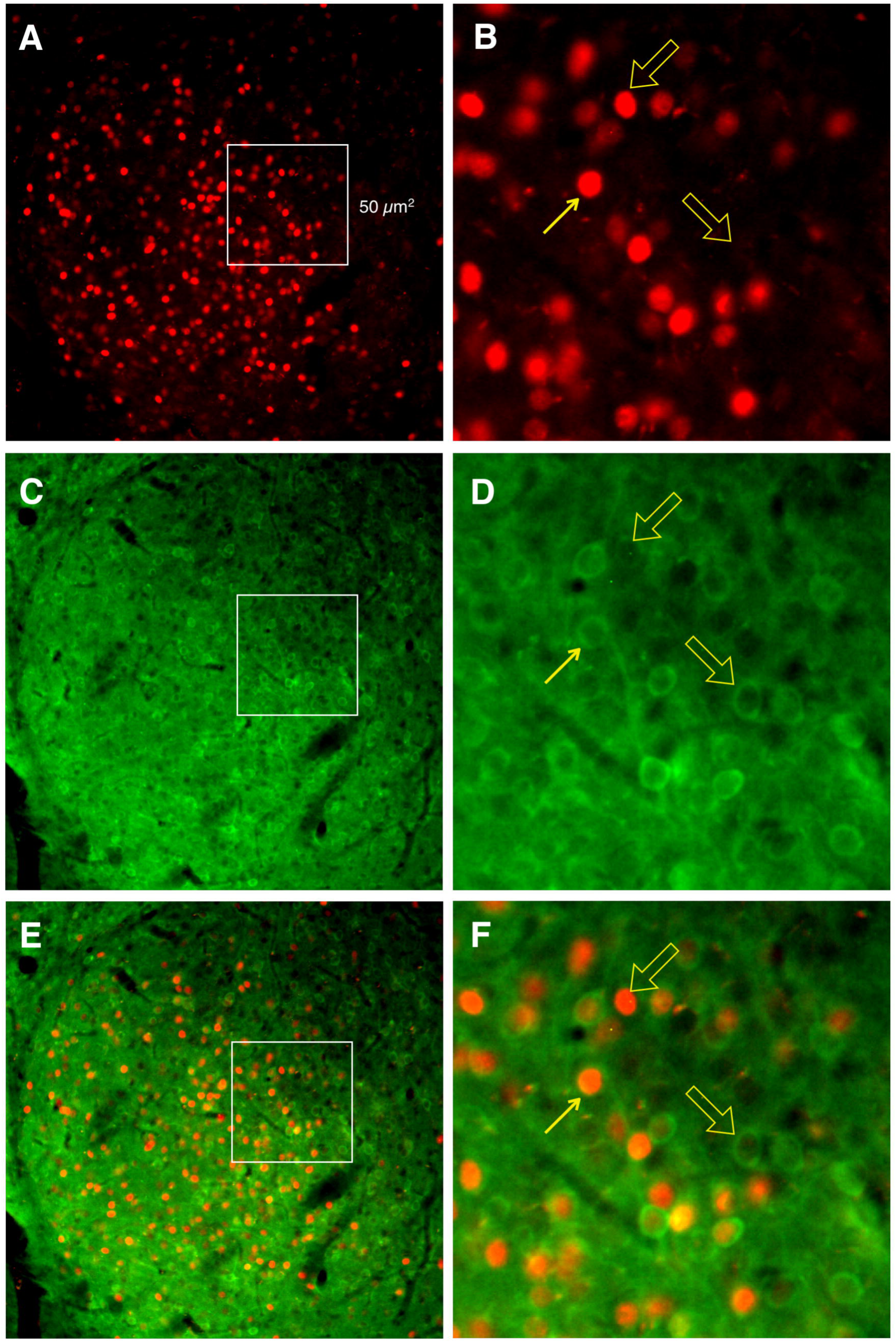
Photomicrographs of fluorescent immunohistochemistry of TORC1 and c-Fos in the CeA 60 min after LiCl injection. Red nuclear c-Fos labeling (A) and magnified (B). Green TORC1 staining (C) and magnified (D). Merged c-Fos and TORC1 labeling (E) and magnified (F). Scale bar 50 µm.

**Figure 10.**
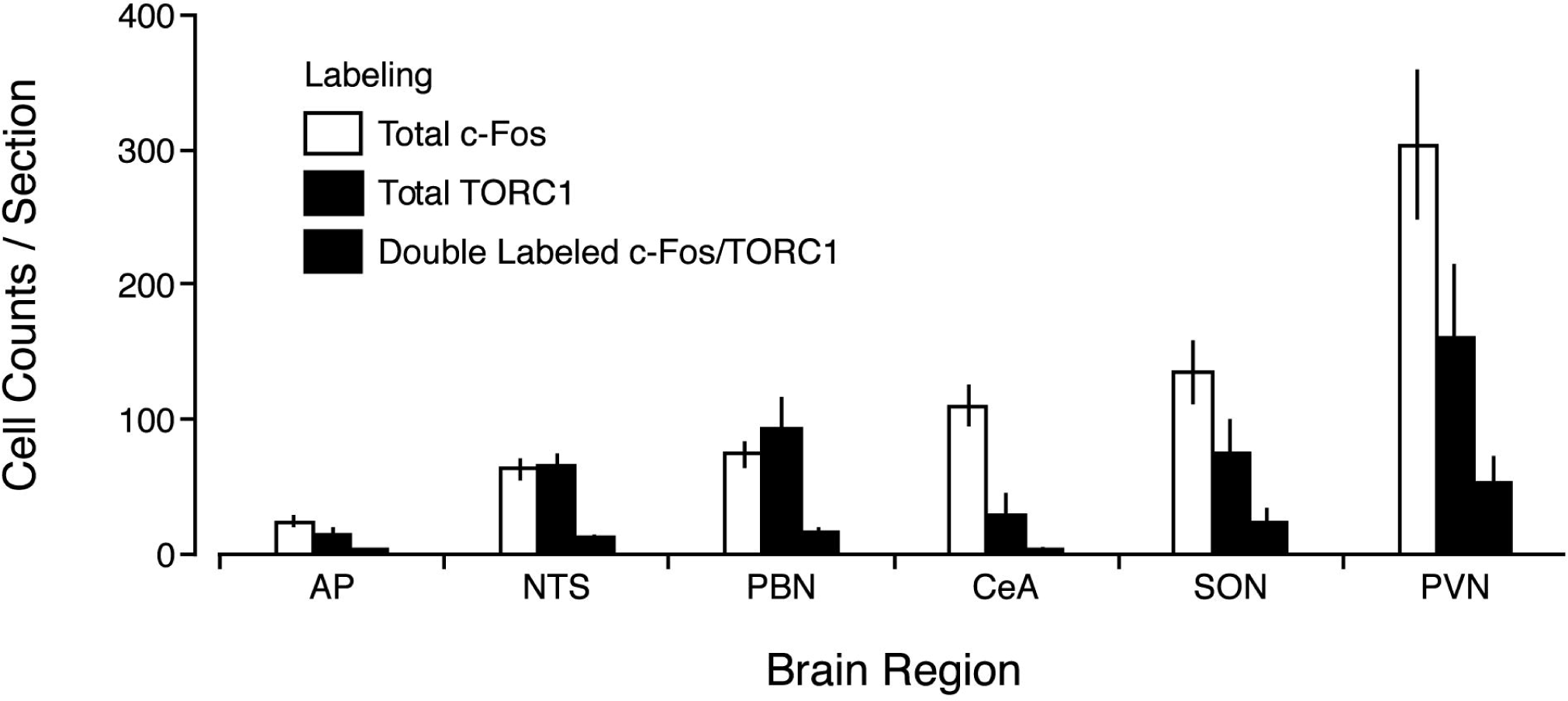
Quantification of nuclear co-localization in the visceral neruaxis between TORC1 and c-Fos 60 min after LiCl injection. Total c-Fos (white bars) and total TORC1 (hatched bars) were significantly greater than the number of double-labeled cells (black bars) in all brain regions.

TORC1 labeling was different depending on the brain region. In the CeA, labeling was either very clearly nuclear (punctate nuclear staining) or distinctly perinuclear and cytoplasmic (easily distinguishable white nuclei surrounded by somatic TORC1 labeling). In other regions (AP, NTS, PBN, PVN, and SON) labeling was either very punctate and nuclear, or diffuse cytoplasmic labeling not directly surrounding the nucleus. In the NTS, many of the neurons expressing nuclear TORC1 ran along the border between the AP and NTS. In the PVN, TORC1 labeling appeared to be mostly in the magnocellular region and almost none was present in the parvocellular portion.

In the AP, an interaction between treatment (LiCl or NaCl) and timepoint (30, 60, 180 min) was found (F(2,30)=3.32, p<.05) for nuclear TORC1. Post-hoc Tukey’s HSD revealed that the 60-min LiCl group had significantly greater nuclear TORC1 than all NaCl groups (30, 60, 180 min) and the 180 min LiCl group. The 30-min LiCl group was not significantly different from any group, suggesting an intermediate effect of LiCl.

In the NTS, an effect of treatment (F(1,30)=4.47, p<.05) was found for nuclear TORC1. No effect of timepoint and no interaction were found. Post-hoc t-test analysis showed that when collapsed into NaCl or LiCl groups, regardless of timepoint, LiCl-induced TORC1 was significantly greater than NaCl-induced TORC1 (p<.05).

In the PBN, no effect of treatment (F(1,30)=3.524, p=.07) or time point, and no interaction were found for nuclear TORC1.

In the SON, no interaction and no effect of treatment or timepoint were found for nuclear TORC1.

In the PVN, an interaction between treatment and timepoint was found (F(2,30)=4.92, p<.05) for nuclear TORC1. Post-hoc Tukey’s HSD revealed that the 60-min LiCl group was significantly greater than all other groups.

In the CeA, an interaction between treatment and timepoint was found (F(2,30)=4.11, p<.05) for nuclear TORC1. Post-hoc Tukey’s HSD revealed that the 30-min and 60-min LiCl groups were significantly <u>lower</u> than the 180-min LiCl group. Qualitatively, TORC1 in the CeA was in most cases either almost all nuclear or almost all cytoplasmic in each animal and appeared to generally be nuclear in NaCl-injected rats regardless of timepoint and in 180 min LiCl-injected rats. TORC1 appeared to be cytoplasmic in 30-min and 60-min LiCl-injected rats. This observation, along with the quantitative analysis, suggests that LiCl causes TORC1 in the CeA to go from a nuclear state to a cytoplasmic state beginning around 30 to 60 min, and returning to a nuclear state by 3 h.

### 2.2 Experiment 2: Nuclear co-localization of TORC1 and c-Fos

We examined nuclear co-localization of TORC1 with c-Fos in the visceral neuraxis at 1 h post-LiCl injection. Each brain region examined showed high levels of c-Fos and TORC1 labeling. Very little double-labeling was noticeable in any region, although some was present.

Similar to the chromogenic labeling, in the CeA TORC1 was found in the cytoplasm after LiCl injection.

In all brain regions (AP, NTS, PBN, PVN, SON, and CeA), both the number of double-labeled cells as a percent of total c-Fos, and the number of double-labeled cells as a percent of total TORC1, were significantly different from 100% (p<.05, t=5.25–44.19).

Thus, TORC1 may not be required for c-Fos induction in the visceral neuraxis. This suggests that CREB is not solely responsible for producing LiCl-induced c-Fos and that other transcription factors are involved.

## 3. Discussion

We examined c-Fos at 60 min and nuclear TORC1 in the visceral neuraxis at 30, 60 and 180 min following i.p. LiCl injection to determine if nuclear localization of TORC1 correlated with c-Fos induction. As expected, injection of LiCl led to consistent induction of c-Fos at 60 min, from a low basal level, across the visceral neuraxis. We saw high basal levels of nuclear TORC1 throughout the visceral neuraxis, suggesting a moderate level of CREB-mediated gene transcription is possible under basal conditions. LiCl injection induced three profiles of nuclear TORC1 (increase, decrease, or no change) in the visceral neuraxis. In the AP, NTS and PVN nuclear TORC1 was increased following LiCl injection, similar to increases in c-Fos. Second, in the PBN and SON, although LiCl injection increased c-Fos, it did not alter nuclear TORC1. Third, in the CeA, LiCl injection <u>reduced</u> nuclear TORC1 compared to NaCl injection; in NaCl-injected rats TORC1 was almost exclusively nuclear, whereas LiCl-injected rats had mostly cytoplasmic TORC1. Thus, in the CeA, LiCl injection had opposing effects on TORC1 (decreased) and c-Fos (increased).

Regardless of whether nuclear TORC1 was regionally increased or decreased in the visceral neuraxis by LiCl injection, at the single cell level nuclear TORC1 and c-Fos were not co-localized. As assessed by immunofluorescent double-labeling, nuclear TORC1 is not present in neurons expressing c-Fos at a time consistent with its involvement in gene transcription.

The lack of cellular co-localization suggests that TORC1- and CREB-mediated gene transcription may not be essential for induction of c-Fos following stimulation of the visceral neuraxis with LiCl. Furthermore, because c-Fos and TORC1 do not co-localize, nuclear TORC1 staining reveals a population of cells, distinct from c-Fos-expressing cells, which also responds to LiCl injection in the visceral neuraxis with an upregulation of CREB signaling.

### 3.1 Differential patterns of TORC1

We saw different levels of nuclear localization throughout the visceral neuraxis. We found increased nuclear TORC1 in the AP, NTS, and PVN following LiCl injection. This is consistent with many other studies, in which TORC shows a rapid translocation to the nucleus after stimulation and a later decline back to baseline during recovery, e.g., after restraint stress in the PVN, ACTH stimulation of rat adrenal cortex in vivo, or stimulation of cultured cortical neurons (Li et al., 2009; Liu et al., 2011; Spiga et al., 2011).

Activation of the AP is consistent with its role in detecting blood-borne toxins like LiCl, and is critical for the effects of systemic injections of LiCl. The AP is critical for the effects of systemic injections of LiCl. LiCl stimulates the visceral neuraxis starting with direct stimulation of the AP by LiCl (Tsukamoto and Adachi, 1994). Additionally, lesions of the AP block behavioral responses (such as lying-on-belly) to LiCl and CTA formation (Bernstein et al., 1992), and AP lesions attenuate c-Fos induction (Spencer et al., 2012).

Direct activation of TORC1 by LiCl may account for increased nuclear TORC1 in the AP. LiCl can act directly on some cells to produce TORC activation (Boer et al., 2007) and can enhance CREB-TORC1 binding in cultured cells bathed in LiCl (Heinrich et al., 2009).

Increases in nuclear TORC1 in the NTS may be due in part to transynaptic input from the AP. The AP projects to the NTS and the two regions are directly adjacent to each other (Lundy and Norgren, 2004). Many of the nuclear TORC1 labeled cells in the NTS ran along the border of the AP where the dendritic fields of NTS cells project into the AP. The NTS is also the first central relay of the visceral vagus nerve, and LiCl may activate the NTS by vagal input (Lundy and Norgren, 2004).

The PVN, as part of the hypothalamic-pituitary-adrenal axis, is activated by stressors (Hsu et al., 2014), including LiCl injections (Spencer and Houpt, 2001). Increased nuclear TORC2 in the parvocellular region of PVN has been found 30 min after restraint stress (Liu et al., 2011), and TORC2 is critical for CREB-mediated CRH gene transcription (Liu et al., 2010). TORC1 has a similar activation profile following stress. However, the nuclear TORC1 was primarily found in the magnocellular region of the PVN, where it may be correlated with the posterior pituitary response to LiCl, such as release of oxytocin and vasopressin (Verbalis et al., 1986).

Interestingly, the CeA showed decreased nuclear TORC1 in response to LiCl injection; in fact, almost all LiCl-treated animals showed only cytoplasmic TORC1. Thus, visceral stimulation appears to cause TORC1 to be transiently exported from the nucleus of CeA cells, in an inversion of the expected stimulation-induction pattern. While removal of TORC1 from the nucleus has been correlated with a rebound decrease in CREB-mediated gene expression during recovery from stimulation, a stimulation-induced decrease of basal nuclear TORC1 is also possible. For example, stimulation of rat liver cells with insulin led to removal of TORC2 from the nucleus and increased cytoplasmic TORC2 (Koo et al., 2005). It has also been suggested that nuclear TORC is initially decreased from baseline levels following ACTH stimulation in adrenal cells, correlated with a transient decrease in CREB-inducible gene expression (Takemori et al., 2003).

The removal of TORC from the nucleus is facilitated by phosphorylation by AMPK family members, particularly salt-inducible kinase (SIK) (Katoh et al., 2006). Thus, the stimulation-induced decrease of nuclear TORC1 in the CeA may be mediated by LiCl-induced SIK activation.

### 3.2 Signaling upstream of TORC1

An increase or decrease in nuclear TORC1 would suggest several things about activity of upstream regulators. Activation of cAMP, PKA, and calcineurin have been shown to lead to dephosphorylation of TORC and its release from protein 14-3-3 sequestration, allowing it to translocate from the cytoplasm to the nucleus (Bittinger et al., 2004; Kingsbury et al., 2007; Li et al., 2009; Screaton et al., 2004; Spencer and Weiser, 2010). TORC is subsequently removed from the nucleus when phosphorylated by AMPK family members (Katoh et al., 2006). Many of these signaling factors have previously been examined, e.g., in CTA learning, which depends on visceral neuraxis stimulation.

Calcineurin appears to act as a constraint on CTA learning: calcineurin activity was decreased in the amygdala 3 days after conditioning, calcineurin knockdown in the mouse forebrain prior to conditioning enhanced learning, and forebrain overexpression decreased learning (Baumgartel et al., 2008).

cAMP and PKA have also been shown to be important for CTA learning. Increasing cAMP in the insular cortex (IC) has been shown to enhance CTA learning (Miranda and McGaugh, 2004), while disruption of PKA in the amygdala has been shown to impair formation of a long-term CTA memory and enhance the rate of CTA extinction (Koh and Bernstein, 2003; Koh et al., 2003).

Additionally, we have found that functional knockout of protein 14-3-3 in the forebrain (including basolateral amygdala and IC) disrupts learning of a CTA (unpublished data).

AMPK family members such as AMPK and SIK have yet to be examined in CTA learning. The findings of TORC1 regulation after LiCl injection suggest the AMPK family may be good candidates for future studies of visceral processing and CTA learning.

### 3.3 CREB may not induce c-Fos

We found that nuclear TORC1 was induced by LiCl, but rarely co-localized in the same cell with c-Fos, suggesting that TORC1 is neither necessary nor sufficient for c-Fos induction in many brain regions. Because TORC is a required co-factor of CREB (Bittinger et al., 2004; Conkright et al., 2003a; Conkright et al., 2003b; Iourgenko et al., 2003) and TORC1 is the most abundant form of TORC in the brain (Watts et al., 2011), the lack of co-localized TORC1 also suggests that CREB itself is not critical for c-Fos induction.

There are several alternative scenarios in which CREB might still contribute to c-Fos expression in a single cell. There are 3 isoforms of TORC, so low-levels of other isoforms may contribute to c-Fos induction without evident cellular co-localization. Alternatively, the sampling times may have missed a transient nuclear localization of TORC1 in cells that ultimately express c-Fos.

### 3.4 Other transcription factors induce c-Fos

However, the lack of TORC1 co-localization suggests the involvement of other transcription factors in c-Fos induction after LiCl injection. In addition to a CRE, the c-Fos promoter contains a serum response element (SRE), a c-Sis inducible element (SIE) and an AP-1/CRE element (FAP). Thus, c-Fos transcription can be enhanced at these sites by serum response factor/ELK-1 complex, or STAT family members, or Fos:Jun dimers, respectively (Herdegen and Leah, 1998; Treisman, 1995). Furthermore, binding at these sites can drive transcription even after deletion of the CRE (Robertson et al., 1995), so they may activate c-Fos in vivo, independent of CREB/TORC. AP-1 family members have been examined previously in CTA learning (Kwon et al., 2008) and other stress-inducing paradigms (Honkaniemi et al., 1992). However, serum response factor and STAT family members have not been explored and provide novel targets to examine after stimulation of the visceral neuraxis with LiCl and during CTA learning.

### 3.5 Conclusions

We hypothesized that TORC1 would be necessary and sufficient for induction of c-Fos in the visceral neuraxis following LiCl stimulation. We also predicted that LiCl would positively regulate nuclear TORC1 in the visceral neuraxis, similar to inductions of c-Fos seen by LiCl and nuclear TORC1 regulation by LiCl would precede c-Fos.

We found that nuclear TORC1 and c-Fos are not present in the same cells in a high percentage of cells. TORC1 and c-Fos are both active separately following LiCl injection, but in parallel. Because the TORC1 cells are a separate population of cells from the c-Fos cells, and LiCl also regulates them, this study identified a new population of cells in the visceral neuraxis that responds to LiCl in parallel to c-Fos-positive cells. This means that TORC1, and thus CREB, is likely not necessary or sufficient for c-Fos induction in the visceral neuraxis following LiCl injection. This result was unexpected, as prior studies have implied a PKA to CREB to c-Fos pathway after LiCl (Houpt et al., 1994; Yasoshima et al., 2006). It should no longer be assumed that LiCl induced c-Fos is the result of CREB-mediated gene transcription. If CREB is not responsible for c-Fos induction then other transcription factors must be involved. Determining which transcription factors produce c-Fos following LiCl stimulation will be necessary for future studies.

## 4. Experimental Procedures

### 4.1 Animals

Adult male Sprague-Dawley rats ranging from 380-650g were individually housed under a 12-h light –12-h dark cycle (lights on 07:00) at 22°C with free access to Purina rodent chow and distilled water. All procedures were conducted in the first half of the lights-on period. Experiments were approved by the Florida State University institutional animal care and use committee.

### 4.2 Tissue collection

At the indicated time after LiCl or NaCl injection, rats were anesthetized with sodium pentobarbital (104 mg/0.4 ml) and perfused first with 100 ml of isotonic saline containing 0.5% sodium azide and 1000 U heparin, and then with 400 ml phosphate-buffered 4% paraformaldehyde. The brains were removed and post-fixed for 24 h, then cryoprotected in 30% sucrose solutions for 1–2 days. The brains were cut at 40 µm on a -20°C microtome.

### 4.3 Immunohistochemistry

Brain sections were washed twice in 0.1 M sodium phosphate-buffered-saline (PBS) for 10 min then permeabilized in 0.2% Triton-1% bovine serum albumin (BSA)-PBS for 30 min. After two PBS-BSA washes, sections were incubated overnight at 25°C with the appropriate primary antibody.

For chromogenic staining, sections were incubated in one of two primary antibodies: anti-c-Fos polyclonal rabbit antisera (Ab-5, Calbiochem, 1:10,000) or anti-TORC-1 monoclonal rabbit antibodies: (C71D11, Cell Signaling, 1:1000). Sections were then washed twice with PBS-BSA for 10 min. Sections were then incubated for 1 h with a biotinylated secondary antibody (goat anti-rabbit antibody, Vector Laboratories, 1:200). Antibody complexes were amplified using the Vectastain ABC Elite kit (Vector Laboratories). Following amplification sections were washed twice with 0.1M phosphate buffer (PB) for 10 min. Immunoreactivity was then visualized by a 5-min reaction with 0.05% 3,3-diaminobenzidine tetrahydrochloride. Sections were then immediately washed twice in 0.1 M PB and mounted on gelatin-coated slides. Slides were coverslipped with Permount.

For fluorescent labeling, sections were co-incubated with anti-c-Fos polyclonal goat antisera (sc-52, Santa Cruz, 1:1000) and anti-TORC-1 monoclonal rabbit antibodies: (C71D11, Cell Signaling, 1:500). After co-incubation of the primary antibodies the sections were washed twice with PBS-BSA for 10 min and incubated for 90 min in the dark with fluorescent secondary antibodies (Alexa 488 anti-rabbit, Cell Signaling, 1:200 and Alexa 555 anti-goat, Invitrogen, 1:200). Sections were washed twice with PBS-BSA for 10 min in the dark and taken to a dark room to mount and coverslip with Vectashield mounting medium (Vector).

### 4.4 Quantification

For chromogenic IHC, cells expressing darkly positive, nuclear staining were quantified using custom software (MindsEye, Thomas A. Houpt). Regions were digitally captured at 40x magnification on a Macintosh computer using an Olympus Provis AX-70 microscope with a Dage-MTI DC-330 CCD camera and Scion LG-3 framegrabber. Counting was restricted to the area delineated by a hand drawn outline. Unilateral cell counts were averaged for sections for each rat. The individual mean counts for each region were averaged across rats within experimental groups. Unilateral cell counts were averaged for 2 sections of the PVN (approximately bregma -1.8 mm to -2.1 mm), 4 sections of the SON (bregma -1.3mm to -1.8mm), 2 sections of the CeA (bregma -2.3 mm to 3.1 mm), 2 sections of the PBN (-8.85mm to -9.5mm), 4 sections of the NTS, (-13.6mm to -13.7mm), 2 sections of the AP (-13.6mm to -13.7mm) (Swanson, 1992).

TORC1 counts were normalized to NaCl controls in order to maintain consistency between timepoints. For TORC1 Two-way ANOVAs between treatment (LiCl or NaCl) and timepoint (30, 60, 180 min) were performed for each brain region (Statistica). Post-hoc Tukey’s Honestly Significant Difference (HSD) for unequal n were performed following ANOVA analysis. c-Fos section counts were not normalized and t-tests were performed between LiCl and NaCl groups for each brain region.

Fluorescent IHC images were captured from the PVN, SON, CeA, PBN, NTS, and AP using the same coordinates listed above at using a Nikon AZ100M microscope and a Nikon Ds-Qi1Mc camera. A single image was taken of each brain region within the coordinates listed above. Sections were hand counted for total TORC1 and c-Fos and double-labeling. One sample t-tests (one-tailed) were run comparing the mean number of double-labeled cells either as a percent of total c-Fos cells vs. 100%, or, as a percent of total TORC1 cells vs. 100%.

### 4.5 Experiment 1: Time course of LiCl-induced nuclear TORC1 and c-Fos expression

#### 4.5.1 Basal Levels of TORC1

To determine the effects of an injection procedure on basal levels of nuclear TORC1, rats were injected with either 0.15M NaCl (12 ml/kg) and perfused 30 min later, or perfused without treatment (n=5–6/group).

#### 4.5.2 TORC1 and c-Fos induction after LiCl Injection

To determine the effects of acute LiCl, rats were injected with either 0.15M LiCl or NaCl (12 ml/kg 0.15M). Rats were then perfused at 30, 60 and 180 min after injection (n=6/group).

Brains were removed, post-fixed, and processed for chromogenic TORC1 immunohistochemistry at 30, 60 and 180 min. In addition, alternate sections from the 60-min LiCl and NaCl groups were processed for c-Fos immunohistochemistry in order to make comparisons between TORC1 and c-Fos induction.

TORC1 and c-Fos positive nuclei were quantified in the AP, NTS, PBN, SON, PVN, and CeA. Because of the high basal numbers of TORC1-positive cells, TORC1 counts at each time point were normalized to the control group for each brain region. Because of the low level of c-Fos in control groups, absolute c-Fos counts are presented.

### 4.6 Experiment 2: Nuclear co-localization of TORC1 and c-Fos

Rats (n=6) were injected with 0.15M LiCl (12 ml/kg) and perfused 1 h after injection. Brains were removed, post-fixed and double-labeled for both c-Fos and TORC1 immunofluorescence. Each rat had one representative section from the AP, NTS, PBN, SON, PVN, and CeA imaged for both c-Fos and TORC1 labeling. Each image was hand counted for single- and double-labeling, and number of co-localized cells. The percent of c-Fos cells co-localized with TORC1 cells and percent of TORC1 cells co-localized with c-Fos cells were both calculated.

## Acknowledgements

Supported by NIDCD T32-000044 (AK).

